# Sonicated fibrils of huntingtin exon-1 preferentially seed neurons and produce toxic assemblies

**DOI:** 10.1101/2021.04.16.440200

**Authors:** Anjalika Chongtham, J Mario Isas, Nitin K Pandey, Anoop Rawat, Jung Hyun Yoo, Tara Mastro, Marry Kennedy, Ralf Langen, Ali Khoshnan

## Abstract

Huntington’s disease (HD) is a genetically inherited neurodegenerative disorder caused by expansion of a polyglutamine (polyQ) repeats in the exon-1 of huntingtin protein (HTT). The expanded polyQ enhances the amyloidogenic propensity of HTT exon 1 (HTTex1), which forms a heterogeneous mixture of assemblies with some being neurotoxic. While predominantly intracellular, monomeric and aggregated mutant HTT species are also present in the cerebrospinal fluids of HD patients, however, their biological properties are not well understood. To explore the role of extracellular mutant HTT in aggregation and toxicity, we investigated the possible uptake and amplification of recombinant HTTex1 assemblies in cell culture models. We found seeding-competent species in the sonicated HTTex1 fibrils, which preferentially entered human neurons and triggered the amplification of neurotoxic assemblies; astrocytes or epithelial cells were not permissive to the HTTex1 seeding. The aggregation of HTTex1 seeds in neurons depleted endogenous HTT protein with non-pathogenic polyQ repeat, activated apoptotic caspase-3 pathway and induced nuclear fragmentation. Using a panel of novel monoclonal antibodies and genetic mutation, we identified epitopes within the N-terminal 17 amino acids and proline-rich domain of HTTex1 mediating neural seeding. Synaptosome preparations from the brains of HD mice also contained similar neurotoxic seeding-competent mutant HTT species. Our findings suggest that amyloidogenic extracellular mutant HTT assemblies may selectively enter neurons, propagate and produce neurotoxic assemblies.

## 1. Introduction

Huntington’s disease (HD) is an autosomal genetically inherited neurodegenerative disorder characterized by debilitating motor, psychiatric, and cognitive symptoms (Bates et al., 2015, Gosh and Tabrizi, 2018). Expansion of a CAG repeat (>35) in the exon 1 of huntingtin HTT gene, which translates into an abnormal polyglutamine (polyQ) tract, is the underlying cause of Huntington’s Disease (HD). (The Huntington’s Disease Collaborative Research Group, 1993). Mutant HTT exon-1 (HTTex1) released by the enzymatic cleavage of full-length protein and/or by aberrant splicing of mutant HTT mRNA, is the most aggregation-prone species, accumulates in the brains of HD patients and is sufficient to induce severe HD-like pathology in animal models (DiFiglia et al., 1997, Davies et al., 1997, Lunkes et al., 2002, Sathasivam, et al., 2010, Bates et al., 2015, Neueder et al., 2018, Yang et al., 2020). The aggregation of mutant HTTex1 produces a heterogeneous mixture of assemblies including oligomers, fibrils and inclusion bodies, which may have different biological properties (Arrasate et al., 2004, Kim et al., 2016, Sahoo et al., 2016, Bäuerlein et al., 2017). For example, transient expression of mutant HTTex1 in culture models revealed that the accumulation of soluble oligomeric species coincide with neurotoxicity (Arrasate et al., 2004, Nucifora et al., 2012). On the other hand, fibrils of HTTex1 may interact with cellular membranes and disrupt their architectures and vital functions (Bäuerlein et al., 2017). The promiscuous interaction of misfolded mutant HTTex1 with cellular proteins may also contribute to the heterogeneity of assemblies and toxicity spectrum (Kim et al., 2016, Wanker et al., 2019).

The oligomerization of mutant HTTex1 *in vitro* occurs by a stepwise mechanism and is influenced by several factors such as interaction with biological membranes and the formation of an oligomeric seed structure, which acts as a nucleating agent and accelerates aggregation (Pandey et al., 2017, Tao et al., 2019). Seed structures may also assemble *in vivo,* however, little is known about their structures and roles in aggregation and toxicity. Seeding-competent mutant HTT assemblies capable of promoting the aggregation of monomeric HTTex1 *in vitro* and in cell models have been isolated from the brains of HD animal models and postmortem brain homogenates and cerebrospinal fluids (CSF) of HD patients. In these studies, the levels of mutant HTT seeds positively correlated with disease severity in the HD patients (Morozova et al., 2015, Ast et al., 2018, Lee et al., 2020). Interestingly, similar to other amyloidogenic proteins such as α-synuclein and Tau, mutant HTTex1 may have the propensity to propagate by a prion-like mechanism (Jucker and Walker, 2018, Pearce and Kopito, 2018, Masnata et al., 2019). For example, mutant HTTex1 assemblies spread between neurons in the brains of *Drosophila* models of HD and form aggregates by recruiting monomeric HTTex1 (Pearce et al, 2018). Moreover, cerebrospinal fluids (CSF) of HD patients seed mammalian cells engineered to express mutant HTTex1-EGFP and induce protein aggregation (Tan et al., 2015, Lee et al., 2020). These encouraging findings provide a new direction in HD research and may have implications for the spread and propagation of neurotoxic mutant HTT assemblies in HD.

The neurodegenerative aspect of HD may release various mutant HTT assemblies in the circulation and CSF. Moreover, HTT is actively and passively secreted from cultured cells and neurons in the HD animal models (Trajkovic, et al., 2017, Caron et al., 2021). Indeed, the levels of mutant HTT in plasma and CSF of HD patient are being used as biomarkers evaluating the efficacy of therapeutics in HD patients (Tabrizi et al., 2019). While few recent studies demonstrate that extracellular amyloidogenic HTTex1 fibrils have the propensity to enter cell lines and neurons in mouse models (Masnata et al., 2019, Lee et al., 2020), the biological properties of extracellular mutant HTT seeds including the mechanism for cell entry, amplification, and neurotoxicity remain to be investigated. Here, we report that some assemblies in the sonicated recombinant HTTex1 fibrils preferentially enter human neurons and produce neurotoxic assemblies by recruiting endogenous HTT with normal polyQ length. Using a panel of novel anti-HTTex1 monoclonal antibodies and mutagenesis, we further show that conformations within the N-terminal 17 amino acids and proline-rich domains (PRD) of HTT are critical for the neural entry and amplification of seeding-competent HTTex1. These studies support the notion that some extracellular assemblies of HTTex1 with tropism for neurons may contribute to the accumulation of toxic assemblies.

## 2. Materials and Methods

### 2. 1. Antibodies

PHP1-PHP3 mouse monoclonal antibodies (mAbs) were reported previously (Ko et al., 2018). The new PHP5 and PHP6 mAbs were isolated from a mAb library made to the N-17 peptide with 7 glutamines at the C-terminus, and PHP7-PHP10 were produced to sonicated mutant HTTex1 fibrils (Khoshnan et al., in preparation). Clones were selected from each hybridoma library for binding to HTT species by ELISA, Western blots and dot blots as previously described ((Ko et al., 2018), (Supplementary figure 2A and B)). Antibody to beta III Tubulin was from Abcam (Cat# ab18207, 1:1000). The secondary antibodies were goat anti-rabbit Alexa Fluor 488 Cat# A32731, goat anti-mouse Alexa Fluor 488, Cat# A28175, goat anti-mouse Alexa Fluor 594 Cat# A32742, Life Technologies, 1:1000 and goat anti-mouse, Horseradish Peroxidase (HRP) Cat# A16072, Invitrogen, 1:10000.

### 2.2. Dot blot assay

A strip of PVDF membrane was pre-wet in 100% methanol for 15 s, soaked in water for 2 min and equilibrated for 5 min in TBS-T (0.05 % Tween, pH 7.4). A sheet of Whatman filter paper was then soaked in TBS-T and placed on a dry sheet of Whatman filter paper on top of some dry paper towels. The PVDF membrane was placed on top of filter stack and 2 μL of each protein was spotted on a pre-marked grid. The membrane was dried to fix proteins to it for 1.5 h at room temperature. The membrane was then blocked, probed with each indicated primary antibody followed by treatment with goat-anti-mouse HRP secondary antibody, and detected by the addition of chemiluminescent agent Clarity^TM^ Western ECL Substrate (Cat# 1705060, Bio-rad).

### 2. 3. Engineering of recombinant PHP2 antibody and lentivirus production

The DNA fragments encoding the antigen binding domain of VH and VL of PHP2 were amplified from a cDNA library made to mRNA of parental hybridoma by standard PCR methods using mixture of commercial primers and sequenced (Khoshnan et al., 2002). The amplified cDNAs were assembled into an IgG2 backbone (provided by Dr. Alejandro Balazs at the Ragon Institute of MGH, MIT and Harvard) by Gibson assembly (New England biolabs) and subsequently cloned into a lentiviral vector (FUGW). Control and PHP2 recombinant viral particles were produced in HEK-293 cells as described previously (Khoshnan and Patterson, 2012). Viral titers were determined using a GFP lentivirus as a reference. Subsequently, MESC2.10 neural progenitor cells (NPCs) were transduced at multiplicity of (2:1). Supernatant of neurons form the transduced NPCs were tested for antibody secretion and PHP2 antibody binding to HTT fibrils was confirmed by Western blots.

### 2.4. Purification of thioredoxin (TRX)-HTTex1 fusion protein

Expression and purification of wild-type (Q25) and mutant (Q46) HTTex1 fusion proteins has been described previously (Bugg et al., 2012, Isas et al., 2015, Isas et al., 2017). Briefly, bacterial cell pellet containing the expressed fusion protein (6XHis-TRXA-HTTex1/or HTTex1-111C variant for labeling) were lysed and cell debris were removed by centrifugation. The clarified lysates containing the fusion protein were loaded onto Ni-column (NiHis60 super flow resin) and washed with a low concentration of imidazole saline buffer. The fusion protein was eluted with high concentration of imidazole saline buffer. The HTTex1-111C variant was first labeled with AlexaFlour 555 (1 to 3 molar ratios for 3hr), diluted 1 to 10 with low salt buffer, loaded on to anionic exchange resin (Mono Q), and further fractionated with a linear gradient (25mM to 1M salt) to remove free label and elute the labelled Alexa sample.

### 2.5. Preparation and purification of HTTex1 fibrils

Monomer HTTex1 (Q25 or Q46) ((HTTex1) (Q46_111Alexa555)) was produced, by the removal of the TRX fusion tag enzymatically (EKMax), followed by separation on reverse phase column (C4) with an acetonitrile gradient as previously described (Pandey et al., 2018). HTTex1 fibrils or (Alexa555 labelled fibrils) were made by first solubilizing lyophilized protein powder from previous step in 0.5% TFA (v/v) in methanol, determined the concentration, and removed the organic solvent by gentle N_2_ gas stream, resulting in a thin clear protein film. The protein film was resuspended in ice cold buffer (20 mM Tris pH7.4, 150 mM NaCl) and adjusted to between 20 to 25 μM concentration. Fibrils reaction started by adding 1% (molar ratio) of sonicated HTTex1 seeds as previously described (Isas et al., 2017) and incubated at 4°C overnight. In a separate reaction, 20% Alexa labelled fibrils were made by adding 4 to 5 μM of HTTex1_111alexa555 solubilized monomer to the Httex1 unlabeled sample. After overnight incubation fibrils were harvested by ultra-centrifugation, and the resulting translucent pellets were resuspended in TFA:H20 (1:4000) at a concentration of 1mg/ml and fragmented using sonication. Mutant HTTex1 in which each residue in PRD domain was replaced with a Pro residue (HTTex1mL17) was generated for probing the binding site of PHP1 and PHP2 as recently reported (Ko et al., 2018). HTTex1mL17 Fibrils were made as described for the HTTex1 in the section above.

### 2. 6. Treatment of cells with HTTex1 seeds

Neurons were derived from embryonic human MESC2.10, IPSC-derived neuronal progenitor cells (NPCs) (provided by Dr. Alysson Muotri, UCSD) according to standard protocols (Khoshnan et al., 2012). Astrocytes were derived from IPSCs by culturing progenitors in DMEM/F12 supplemented with N2 growth factor and 0.5% FBS (Cat# ES-009-B, Millipore) for 14-20 days. Caco-2 and Neuro2A were obtained from ATCC and cultured according to instructions provided. Each line was treated with 10 nM of sonicated HTTx1 fibrils for the indicated time points in the figure legends and processed for immunocytochemistry and confocal microscopy as described previously (Khoshnan, et al., 2012). For antibody inhibition assay, 10 nM of sonicated fibrils were pre-incubated with 1 μg of each antibody clone indicated in the figures for 2 hr at RT and subsequently added to growth medium and processed as above.

### 2.7. In vitro seeding assay with HTT species

Sonicated HTTx1 seeds (10 ng) were incubated with recombinant WT HTTex1 monomers or 100 μg of total protein from human neural lysates for 4 hr at 25°C with continuous agitation. Semi-denaturing detergent agarose gel electrophoresis (SDD-AGE) was performed to examine the amplified products as described previously (Halfman and Lindquist, 2008) with some modifications. Briefly, seeded lysates were loaded in a 1.5% agarose gel in 1X Tris-acetate-EDTA (TAE) (Tris base, glacial acetic acid and EDTA) containing a final concentration of 0.1% SDS. Using 1X Tris-buffered saline (TBS), downward capillary action was used to transfer protein to immune-blot PVDF membrane (Merck cat# IPVH00010). Membranes were blocked with 5% milk in wash buffer (0.05% Tween in PBS) and incubated with the indicated primary anti-HTT antibodies overnight at 4°C. HRP-conjugated goat anti-mouse secondary antibody (1:10000) in wash buffer was then applied for 2 h and developed with ECL substrate and X-ray film.

### 2. 8. Western blot analysis of seeded neurons

MESC2.10 derived neurons grown in 10 cm culture plates were treated with 10 nM of sonicated HTTex1 fibrils for 24 hr. The seeded neurons were harvested and lysed in RIPA buffer (50 mM Tris-HCl (pH 7.4), 150 mM NaCl, 1% NP−40, 0.1% SDS and protease inhibitor). 50 μg of neuronal lysate and corresponding mHTTex1 seed were boiled for 5 min with sample loading buffer and loaded into a 4-20 % polyacrylamide gel (SDS-PAGE, Criterion Bio-Rad) to detect monomeric endogenous HTT or by semi-denaturing detergent agarose (1.5%) gel electrophoresis (SDD-AGE) to probe for the seeded aggregated products. Different antibodies indicated in the figures and figure legends were used for detecting different species of HTT.

For proteinase K resistance assay, 0.1 μg of mHTTex1 seed or 50 μg of seeded neuronal lysate was incubated with varying doses of proteinase K (0 to 0.5 μg/mL, Qiagen) for 30 min at 25°C and heat inactivated at 75°C for 10 min. The digested products were analyzed by SDD-AGE and western blot. PHP2 was used as the detection antibody.

### 2.9. Immunodepletion of HTT in neural lysates

Immunodepletion of HTT was performed by incubating100 μg of neuronal lysates with 2 μg of each anti-HTTex1 or control antibodies overnight at 4°C with rotation. The antibody-HTT complexes were precipitated with protein G covalently conjugated to magnetic beads (Thermo Scientific, P188803). The HTT-depleted lysates were seeded with 10 ng of sonicated HTTex1 fibrils for 4 hr at 25°C with rotation. SDD-AGE was performed to detect the amplified products. The primary antibody used for western blot analysis was PHP1, which is reactive to mutant HTT aggregates in HD animal models (Ko et al., 2018).

### 2.10. Sucrose fractionation of HD brains

Synaptosome and ER/Golgi fractions were prepared from individual mouse forebrains according a recently published method (Mastro et al., 2020). Briefly, forebrains of 9-month old ZQ175 HD mice were dissected from each animal, rinsed in Buffer A (0.32 M sucrose, 1 mM NaHCO_3_, 1 mM MgCl_2_, 0.5 mM CaCl_2_, 0.1 mM phenylmethylsulphonyl chloride (PMSF, Sigma Millipore, St. Louis, MO). Each individual forebrain was homogenized in Buffer A (10% w/v, 4.5 ml for mice) with 12 up and down strokes of a Teflon/glass homogenizer at 900 rpm. Homogenates were subjected to centrifugation at 1400g for 10 min. The pellet was resuspended in Buffer A to 10% w/v (3.8 ml), homogenized (three strokes at 900 rpm) and subjected to centrifugation at 710 g for 10 min. The final resultant pellet (P1) was harvested as a crude fraction containing the nuclei. The two supernatants (S1) were combined and subjected to centrifugation at 13,800g for 10 min. The resulting pellet (P2) was resuspended in Buffer B (0.32 M sucrose, 1 mM NaHCO_3_; 2 ml for mice), homogenized with 6 strokes at 900 rpm in a Teflon/glass homogenizer, and layered onto a discontinuous sucrose gradient (equal parts 0.32M, 0.85 M, 1.0 M, and 1.2 M sucrose in 1 mM NaH_2_CO_3_ buffer (10.5 ml). Gradients were subjected to centrifugation for 2 hours at 82,500g in a swinging bucket rotor. The bands between 0.32M and 0.85M sucrose (light membranes, Myelin), 0.85M and 1.0M sucrose (light membranes, Endoplasmic Reticulum, Golgi), 1.0M and 1.2M sucrose sections (Synaptosomes).

### 2.11. Assessment of seeding activity of protease-resistant brain fractions

Synaptosome and ER/Golgi fractions isolated from ZQ175 mouse brain by sucrose fractionation were dialyzed using Slide-A-Lyzer Dialysis Cassette (Thermo Fisher Scientific, 66203). The dialyzed mouse brain fractions isolated by sucrose fractionation were incubated with 0.01 μg/mL of Proteinase K at 37°C for 1 hr and heat inactivated at 75°C for 10 min. MESC2.10 neurons in 12-well plate were then incubated with 50 μg of each fraction at 37°C for 24 hr. This was followed by immunocytochemistry to examine for the presence of seeding-competent fibrils and assembly formation.

### 2.12. Immunocytochemistry and nuclear damage quantifications

Cells were fixed with 4% formaldehyde in PBS at room temperature for 30 min. After permeabilization and blocking (70% methanol in PBS, at least 1h at −20°C, 10% normal goat serum and 2% BSA in PBS, 30 min at room temperature); cells were incubated overnight with specific primary antibodies at 4°C. After washing in PBS, cells were incubated with appropriate AF 594 or 488 conjugated secondary antibodies in blocking solution for 2 hr at RT, washed and mounted in Vectashield with DAPI. At least 6 random pictures were captured using a Leica SP8 confocal microscope and analyzed using LAS X software. For each picture, the total number of intact cells as well as those with fragmented nuclei were counted. The percent of neurons with intracellular HTT aggregates (seeded neurons) and neurons with nuclear fragmentation was then calculated for each condition and plotted.

### 2.13. Caspase-3 Assays

Neurons were treated with 10 nM of sonicated HTTex1 seeds for 24 hr. PHP1 antibody (1:1000) was used to detect intracellular HTT fibrils and caspase-3 activity in untreated and treated neurons was monitored by immunocytochemistry using anti-caspase-3 antibody (Abcam, Cat# 32351, 1:1000). Images were captured using a Leica SP8 confocal microscope and analyzed by LAS X software. Neurons with aggregates and active caspase-3 were quantified as described in the section above.

### 2.14. Statistical analysis

For pairwise comparisons, Student’s t-test was used and analysis of variance (ANOVA), followed by the Tukey’s post hoc test was performed for multiple comparisons. P values <0.05 were considered as significant and for all results, values are presented as the mean and error bars indicate standard error of the mean (SEM). Statistical significance was assigned as * = P < 0.05, ** = P < 0.01, *** = P < 0.001.

## 3. Results

### 3.1. HTTex1 assemblies preferentially enter human neurons and amplify

To explore the seeding properties of recombinant HTTex1 (46Qs), we incubated pre-assembled fibrils (Isas et al., 2017) with human neurons and examined for uptake and amplification. Mutant HTTex1 fibrils bound to neural surfaces but were not taken up. However, sonication of the fibrils generated smaller species (HTTex1 seeds), which seeded neurons derived from an embryonic neuronal progenitor cell line (MESC2.10) and formed assemblies detected by antibodies specific to the proline-rich domain (PRD) of HTTex1 (Fig. 1A, B). To verify the amplification of HTTex1 seeds, we examined equivalent ratio of the seeds and lysates of seeded neurons by semi-denaturing detergent agarose gel electrophoresis (SDD-AGE) followed by Western Blots (WBs) and found a heterogeneous mixture of assemblies larger than ~ 250 kDa reactive to anti-PRD antibodies (Fig. 1C). The neural seeding and amplification of HTTex1 was progressive and rapid affecting ~60% of the cells by 24 hr (Fig. 1B). Similar to the HTTex1 seeds, the amplified products in neuronal lysates displayed partial resistance to proteinase-K (PK) digestion indicating the formation of complex assemblies (Fig. 1D and E, respectively). Sonicated fibrils of WT HTTex1 (25Q) also entered neurons and amplified suggesting polyQ expansion is not essential for the seeding properties of HTTex1 assemblies (Supplementary fig. 1A). Collectively, these results suggest that seeding-competent HTTex1 assemblies in the sonicated HTTex1 fibrils may preferentially enter neurons and amplify producing a heterogeneous mixture of aggregates.

**Figure 1.**
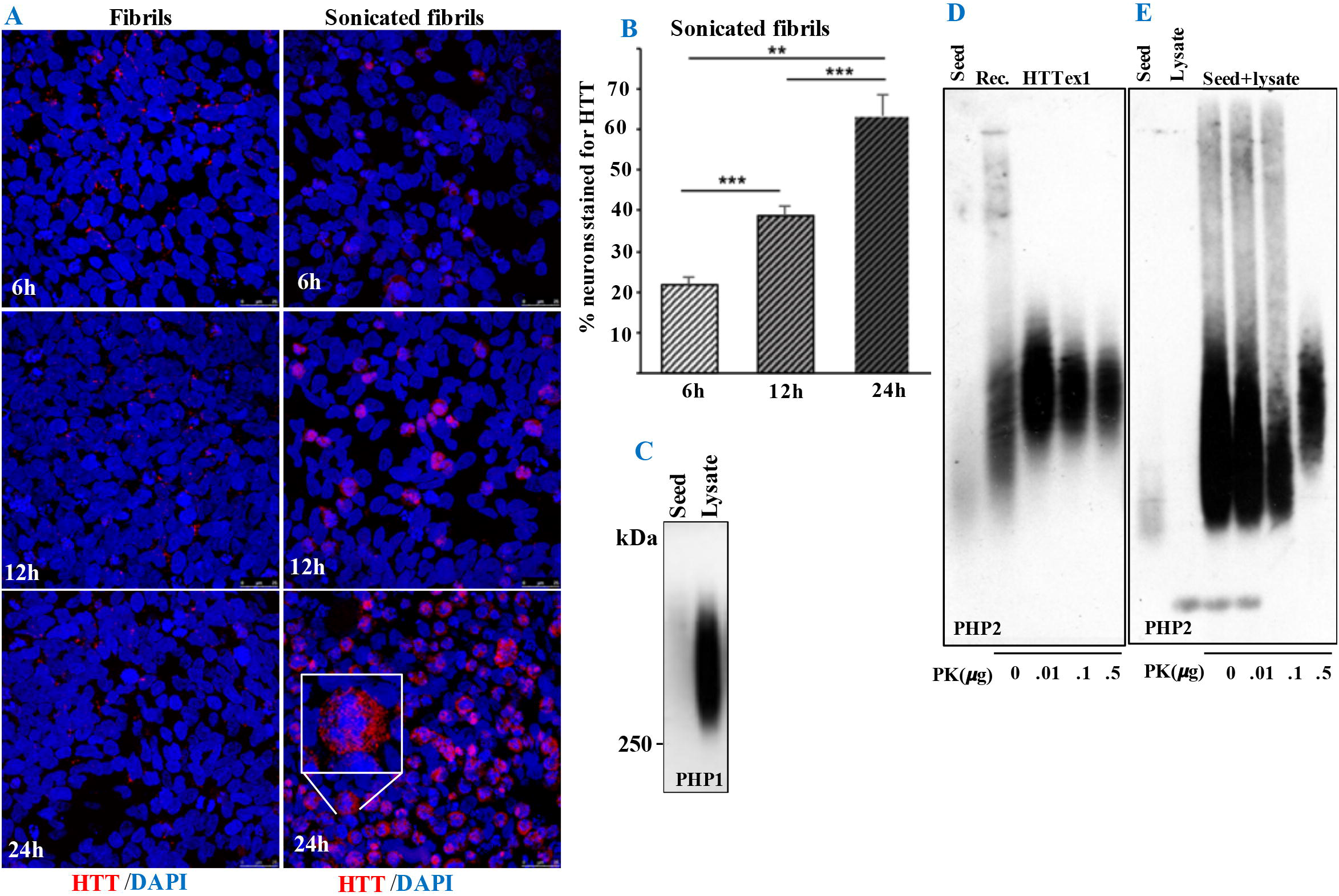
HTTex1 species enter and amplify in neurons. (A) Human neurons derived from MESC2.10 embryonic stem cell line were treated with mutant HTTex1 fibrils or sonicated fibrils (seeds), incubated for the indicated time points and processed for ICC using anti-PRD PHP1 antibody (HTT). Insert in the bottom right panel is a magnified seeded neuron. (B) Quantification of seeded neurons at different time points. (C). Examination of the lysates of seeded neurons 24 hr post-treatment by SDD-AGE detected by PHP1. HTTex1 seed were adjusted to the amounts in the lysates loaded to determine the extent of amplification. (D &E) HTTx1 seeds and its amplified products in MESC 2.10 derived neurons are partially resistant to proteinase K (PK) digestion. Seeds (D) and neural lysates from seeded neurons (E) were treated with an increasing concentration of PK and examined by SDD-AGE and WBs using PHP2 antibody. Data are mean ± SEM; **P<0.01, ***P<0.001.

To determine whether the seeds enter neurons directly, we covalently conjugated the seeds with Alexa 555 fluorophore and added to neurons. By 6 hr post-treatment, labeled HTTex1 predominantly accumulated in the nuclei and increased by longer incubation (Fig. 2A, left panels and B). Notably, the number of neurons with detectable labeled HTTex1 seeds were maximal by ~12 hr of incubation but staining of the seeded neurons by the anti-HTT antibody showed ~ 4-fold increase in reactivity by 24 hr post treatment (Fig. 2A, right panels and B). One possibility is that undetectable levels of the labeled HTTx1 seeds may enter some neurons and promote aggregation, alternatively the newly formed assemblies may spread to the neighboring neurons and propagate. Interestingly, although the HTTex1 seeds entered neurons derived from induced pluripotent stem cells (IPSCs), they did not affect astrocytes produced from the same line even after incubation for up to 4 days in culture (Fig. 2C and D, respectively). We also examined human epithelial cell lines such as Caco-2 or mouse neuro2A cell line but found no evidence for entry and aggregation (Supplementary fig. 1C and D, respectively). These results suggest that seeding-competent HTTex1 assemblies may preferentially enter neurons and accumulate in the nucleus.

**Figure 2.**
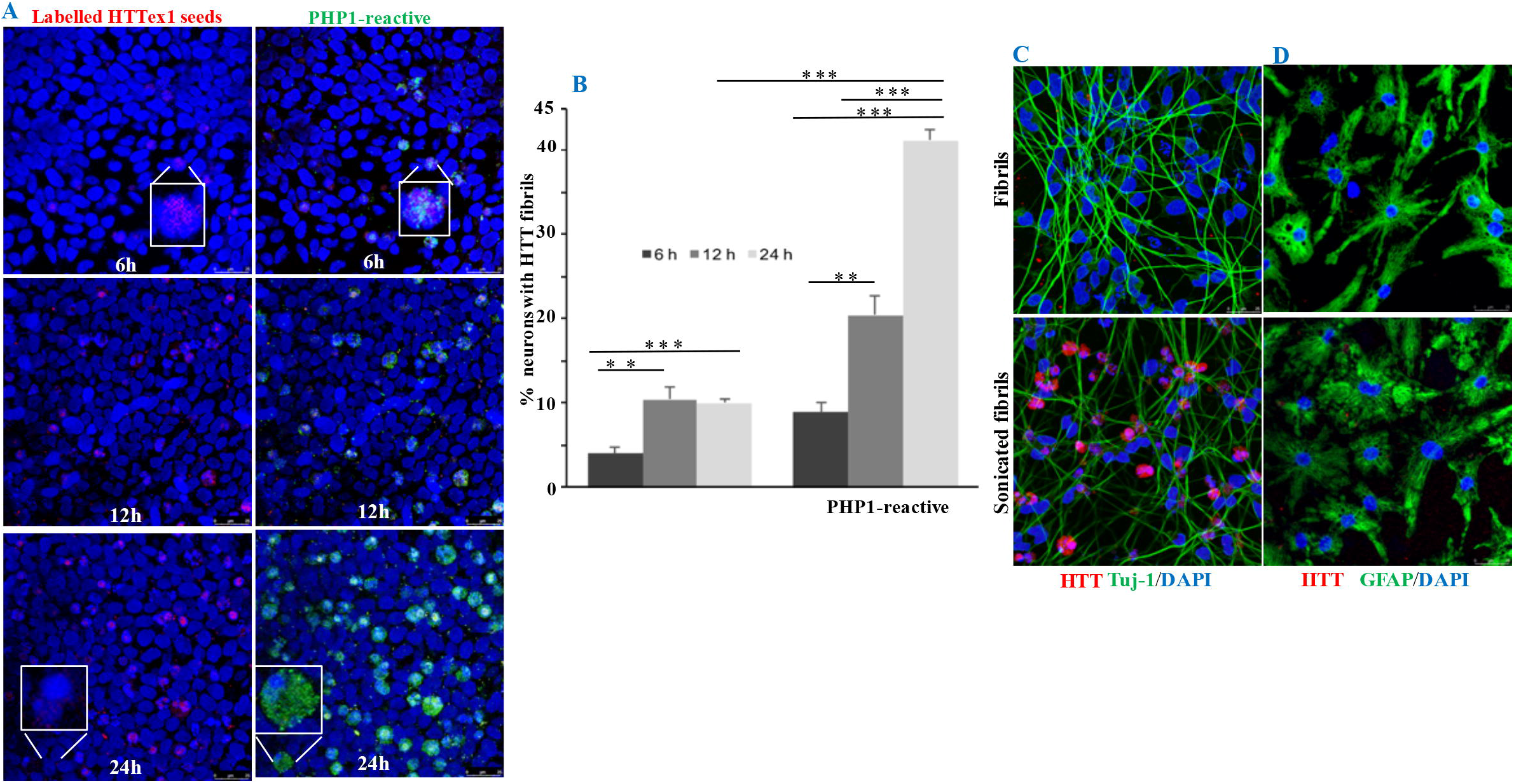
HTTex1 seeds accumulate in the nuclei of neurons. (A) Fluorophore-labelled (Alexa 555) HTTex1 seeds were added to human neurons for the indicated time points and processed for ICC using PHP1 antibody. Confocal images in the left panels show the nuclear localization of labelled HTTex1 seeds and in the right panels are merged images of labelled HTTex1 and amplified species detected by PHP1 antibody. Inserts in the top and bottom panels are magnified neurons. (B) is the quantification of neurons with the labelled HTTex1 (left columns) and neurons labelled with PHP1 antibody (right columns). (C&D) Human neurons from IPSC-derived neuronal progenitors (left panels) or IPSC-derived astrocytes (right panels) were treated with HTTex1 fibrils or sonicated fibrils for 24 hr and processed similar to part (A). Neurons were stained with PHP1 and anti-Tuj-1 and astrocytes were stained with PHP1 and anti-GFAP. Data show mean ± SEM; **P<0.01, ***P<0.001.

### 3.2. Mutant HTTex1 seeds produce neurotoxic assemblies

Importantly, HTTex1 seeds-triggered aggregation promoted robust and progressive nuclear fragmentation, which could be visualized and quantified in the DAPI- stained images (Fig. 3A and B). To confirm the toxicity of HTTex1 seeds, we stained treated neurons for caspase-3 activation and found it activated in the majority of neurons with aggregates (Fig. 3C-E). These findings indicate that the amplification of extracellular HTTex1 seeds in neurons produces toxic assemblies and may constitute a pathogenic aspect of mutant HTTex1.

**Figure 3.**
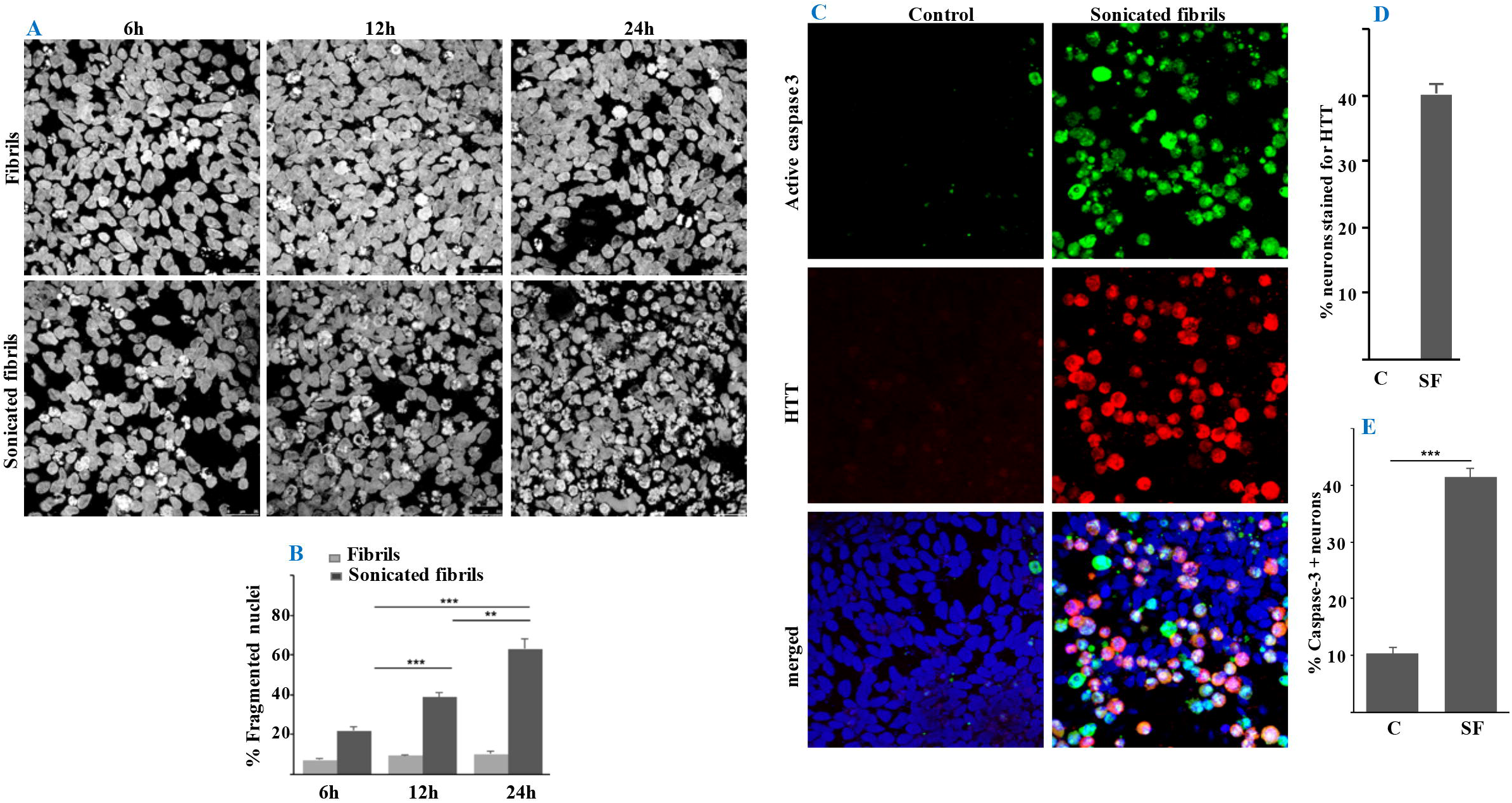
HTTex1 assemblies cause nuclear damage and caspase-3 activation. (A) For a better contrast, black and white images of DAPI stained neurons treated with HTTex1 fibrils or sonicated fibrils for the indicated time points are shown to display the progressive nuclear damage. (B) Quantification of DNA damage in neurons seeded with unsonicated or sonicated mutant HTTex1 fibrils. (C) Confocal images of neurons with active caspase-3, which was induced in neurons treated with the sonicated HTTex1 seeds. Graphs in D and E show quantification of neurons with HTT aggregates and active caspase-3, respectively. Data are expressed as mean ± SEM; ***P<0.001.

### 3.3. Neural entry of HTTex1 seeds deplete endogenous HTT

To better characterize the mutant HTTex1-induced assembly formation in neurons, we examined the lysates of seeded neurons by SDD-AGE followed by WBs using a panel of antibodies reactive to the PRD of HTTex1 seeds (Supplementary fig. 2A and 2B). High molecular weight assemblies (>250 kDa) were detected by several anti-HTT monoclonal antibodies (PHP1, PHP2, PHP7, PHP8), whereas two clones (PHP9 and PHP10) reacted poorly (Figs. 1E, 4A). A likely possibility is that the conformations for the binding of PHP9 and PHP10 may be altered and/or masked in the amplified products. Notably, examination of the seeded neural lysates with antibodies binding to monomeric HTT species (PHP1, PHP5, 2166) revealed depletion of endogenous full-length and N-terminal fragments of HTT (Fig. 4B, arrows). This raised the question of whether the HTTex1 seeds may recruit the endogenous HTT as a substrate. Sequencing the cDNA of HTT in MESC2.10 progenitors revealed 2 copies with 24Qs and 25Qs (data not shown). To confirm that HTTex1 seeds can recruit HTT fragments with normal polyQ length, we developed an *in vitro* amplification assay using monomeric recombinant WT HTTex1 (25Qs) as a substrate. Examined by SDD-AGE, we found that the seeds recruited monomeric HTTex1 and produced assemblies, which migrated similar to those formed in neurons (Figs. 4C and 4A, respectively). Interestingly, HTTex1 seeds also triggered assembly formation in the neural lysates (Fig. 4D, the lysates with several anti-HTT antibodies before adding the seeds. Depletion with the anti-N17 antibodies (PHP5 and PHP6) or clones reactive to the PRD of HTTex1 (PHP1, PHP2 and PHP10) (Supplementary fig. 2A and B) significantly reduced protein aggregation triggered by the HTTex1 seeds (Figure. 4D, Supplementary fig. 2C). Depletion with other anti-HTT antibodies such as PHP3, which binds to polyQ/polyP junction of HTT (Ko et al., 2018) or PHP8 (Supplementary fig. 2A) did not affect the mutant HTTex1-induced aggregation (Supplementary fig. 2C). These studies suggest that HTTex1 seeds may recruit the N-terminal fragments of HTT to propagate in neurons. Given that the 2166 antibody, which binds to epitope outside exon1, does not react with the amplified products (Fig. 4B), it is difficult to determine whether the HTTex1 seeds also recruit full-length HTT. However, reduction in the levels of the microtubule-associated protein Tuj-1 indicates that HTTex1 seeds may recruit neuronal proteins which may include full-length HTT and those interacting with it (Fig. 4B, bottom panels).

**Figure 4.**
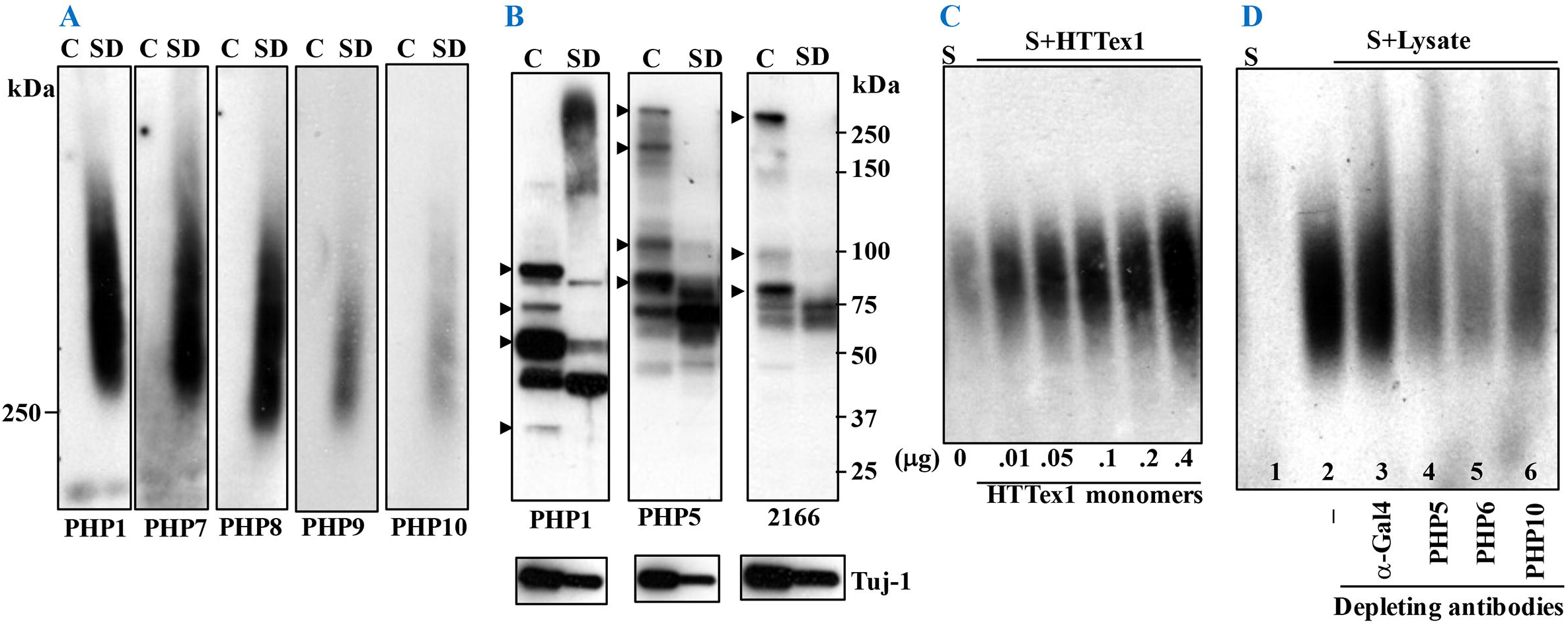
HTTex1 seeds recruit endogenous HTT species to form assemblies. (A) SDD-AGE analysis followed by WBs of control neuronal lysates (C) or lysates of seeded neurons (SD) with sonicated HTTex1 fibrils probed with the indicated anti-HTT antibodies. Part B is SDS-PAGE of similar lysates probed with antibodies reactive to soluble HTT. Arrowheads indicate depletion of endogenous full-length and N-terminal fragments of HTT in the lysates of seeded neurons. (C) Seeding of recombinant WT HTTex1 monomers by the HTTex1 seeds (S). Products were examined by SDD-AGE and WB using PHP1 antibody. (D) Neural lysates were seeded with the mutant HTTex1 seeds. Lane 1 is seed alone and lane 2 is seeded neuronal lysate. The indicated antibodies below each lane were used to deplete the endogenous HTT species before seeding. The binding epitopes of PHP antibodies are shown in supplementary fig. 2A.

### 3.4. Antibodies to conformations in the N17 and PRD block neural entry and amplification of HTTex1 seeds

To identify the epitopes regulating the neural entry and propagation of HTTex1, we incubated the HTTex1 seeds with antibodies reactive to various conformations (Supplementary fig. 2A) and added the seed-antibody complexes to neuronal cultures. We found that pre-incubation of HTTex1 seeds with antibodies to the N17 domain (PHP5, PHP6) and clones reactive to the PRD (PHP1, PHP2, and PHP10) blocked neural entry, amplification and nuclear damage. Importantly, two anti-PRD antibodies PHP7-PHP9, which also bind the seeds (Supplementary fig. 2B), were ineffective suggesting that distinct conformations or regions in the PRD of HTTex1 seeds may be critical for entry (Fig. 5A and B, Supplementary fig. 3). HTTex1 seeds pre-incubated with the N17 antibodies (PHP5, PHP6) remained bound to neurons but did not enter. One possibility is that HTTex1 interacts with neurons using epitopes within the PRD, however, antibodies binding to N17 of HTTex1 seeds may prevent entry by affecting yet unidentified step. To verify that the epitope in the PRD is critical for the neural entry and amplification of HTTex1, we mutated the known binding site of the protective antibodies PHP1 and PHP2 (L17 domain, Supplementary fig. 2A), (Ko et al., 2018). Consistent with the antibody inhibition assays, sonicated fibrils of the PRD-mutated HTTex1 (HTTex1mL17) were incapable of seeding neurons and inducing nuclear damage (Fig. 5C and D). Collectively, these results suggest that specific conformations in the N17 and PRD are critical for the neural entry and propagation of HTTex1 seeds.

**Figure 5.**
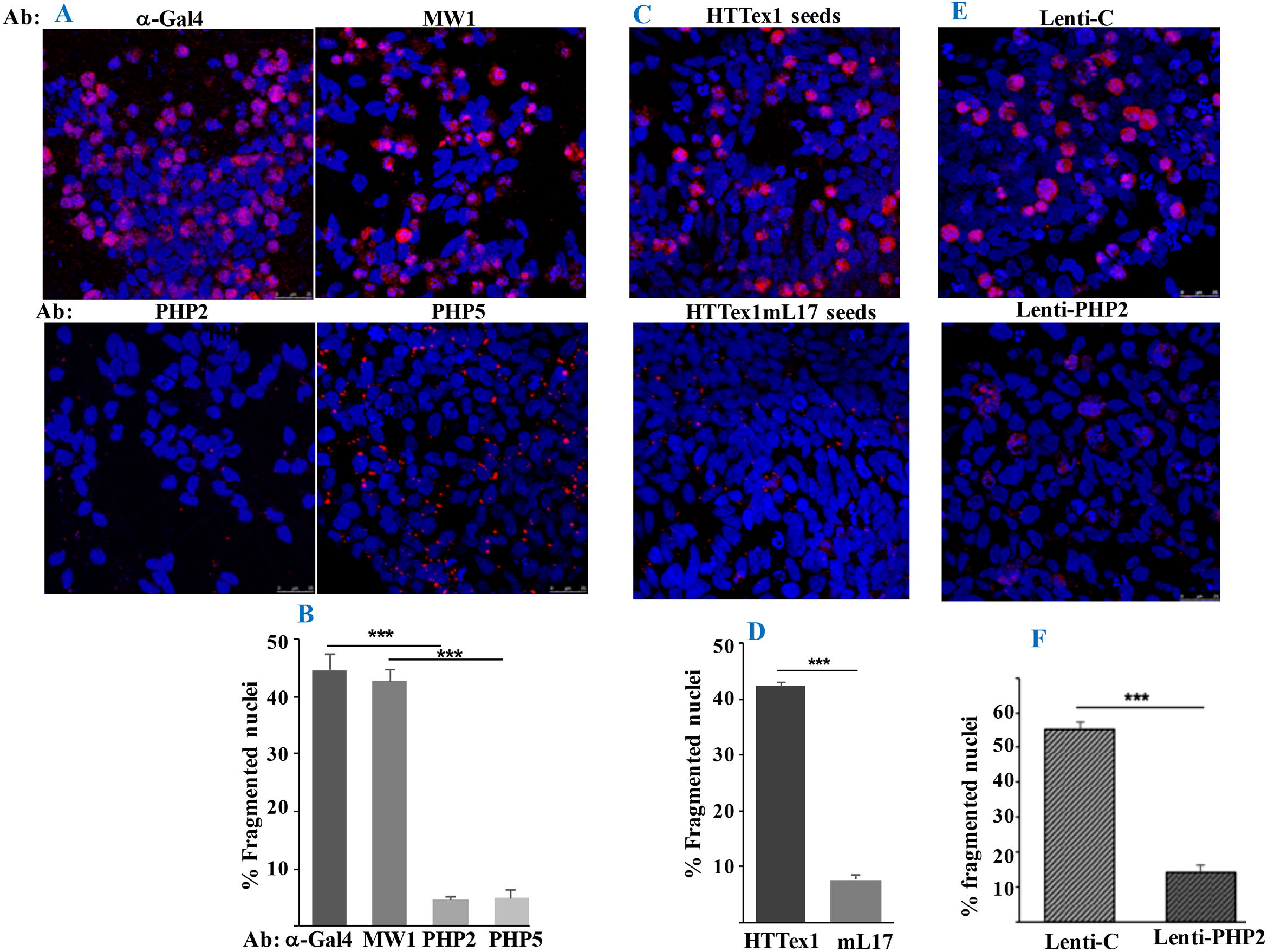
Antibodies to N17 and PRD of HTTex1 block neural seeding. (A) HTTex1 seeds were pre-incubated with the antibodies indicated on top of each panel, subsequently added to neurons and incubated for 24 hr. Samples were processed for ICC, stained with PHP1 and examined by a confocal microscope. Panel B is the quantification of fragmented nuclei in each condition. (C) HTTex1 seeds or those generated from HTTex1 with mutated PRD (HTTx1mL17) were added to neurons and processed as in A. Anti-N17 antibody PHP5 was used to detect the amplified products. Panel D is the quantification of fragmented nuclei in each condition. (E) MESC2.10 neurons engineered to secret control or PHP2 recombinant antibodies (Lenti-PHP2) were treated with the HTTex1 seeds, incubated for 24 hr, processed for ICC using PHP1 antibody and examined by a confocal microscope. Panel F is the quantification of fragmented nuclei in each condition. Data show mean ± SEM; ***P<0.001.

### 3.5. Neurons secreting anti-HTT antibodies are immune to HTTex1 seeding

To explore the therapeutic application of antibodies preventing the neural entry of extracellular HTTex1, we engineered neurons to secrete PHP2 antibodies. MESC2.10 NPCs were transduced with a lentivirus encoding full-length PHP2 antibody. Secreted antibodies from neurons react with the HTTex1 seeds (Supplementary fig. 4). We found that neurons secreting the recombinant PHP2 are resistant to seeding by the HTTex1 since the entry, amplification and nuclear damage were significantly reduced (Fig. 5E and F). These data further confirm the inhibitory properties of PHP2 and support the potential use of recombinant antibodies to prevent seeding by extracellular HTTex1 assemblies in HD models.

### 3. 6. Seeding-competent mutant HTT species accumulate in the brains of ZQ175 HD mice

To determine whether seeding-competent mutant HTT species accumulate *in vivo*, brain homogenates of 9-month old ZQ175 HD mice were centrifuged on sucrose gradients optimized to enrich for myelin, ER/Golgi and synaptosome fractions (Mastro et al., 2020) and examined for the presence of mutant HTT by SDD-AGE and WB (Figure. 6A). Aliquots of ER/Golgi and synaptosome fractions, which had the most HTT species, were subsequently examined for entry and propagation in neurons. We found mutant HTT assemblies predominantly in the synaptosome fractions of some animals were capable of seeding neurons (Fig. 6B), however, the seeding activity was not consistently present in the preparations from various cohorts of similar aged animals (Fig. 6C, top panels). We predicted that similar to antibody inhibition assays (Fig. 5A), HTT-interacting proteins may interfere with the neuronal entry of mutant HTT. Since HTTex1 seeds and those amplified in neurons are resistant to partial proteinase K (PK) digestion (Fig. 1D and E), we performed partial PK digestion of ER/Golgi and synaptosome fractions prior to testing in neurons. Interestingly, PK treatment significantly enhanced the seeding activity of mutant HTT in the synaptosome and to some extent in the ER/Golgi fractions (Fig. 6C, bottom panels). We predict that PK treatment may expose the critical conformation of mutant HTT seeds by digesting the interacting proteins; alternatively, PK treatment may directly produce seeding-competent species by cleaving mutant HTT. Similar to recombinant HTTex1, seeding by the mutant HTT assemblies from HD animals induced nuclear fragmentation (Fig. 6D). These data suggest that seeding-competent mutant HTT species may accumulate in the brains of HD animals and are consistent with the recent findings where the mutant HTT isolated from the brains of ZQ175 HD mice seeded HEK-293 cells expressing mutant HTTex1-EGFP (Lee et al., 2020).

**Figure 6.**
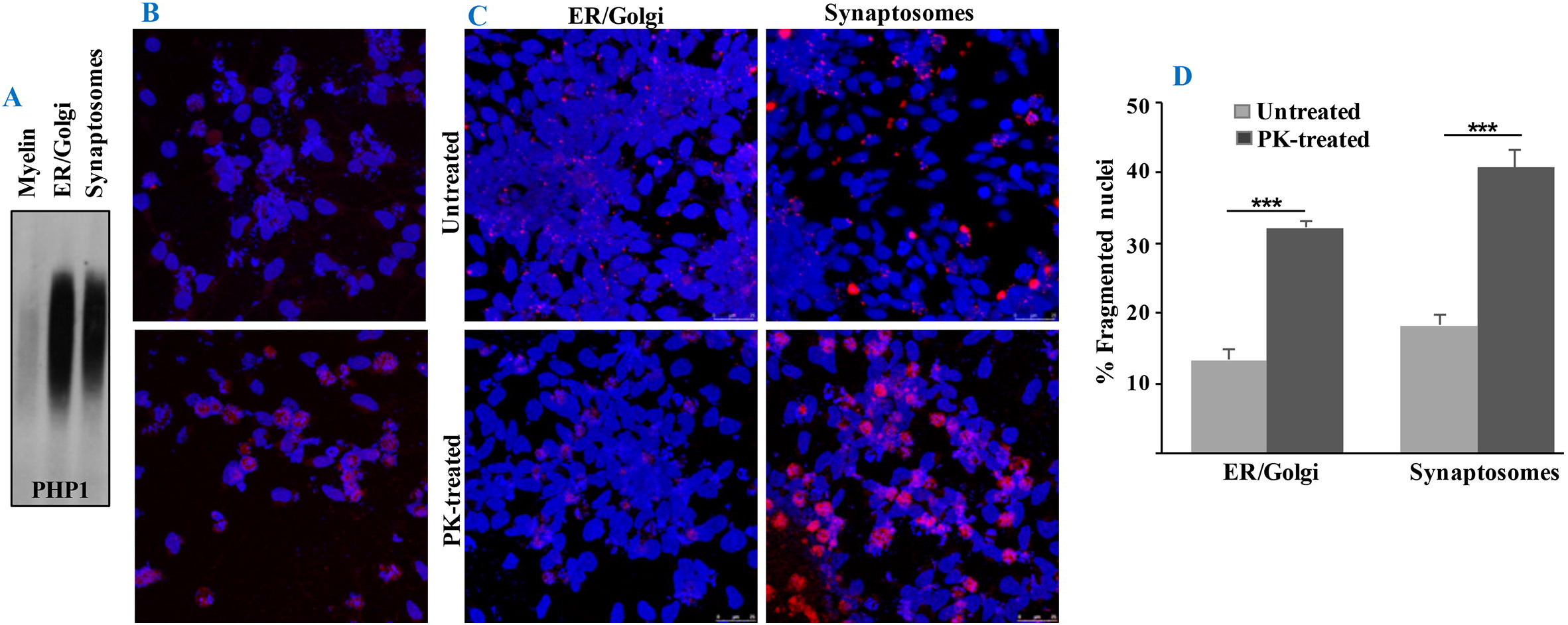
Mutant HTT seeds accumulate in the brains of ZQ175 HD mice. (A) Fractionation of mouse brains on sucrose gradients and examination by SDD-AGE and WB. HTT was detected by PHP1 antibody. (B) Dialyzed aliquots of each fraction (50 μg) were sonicated and added into human neurons derived from MESC2.10 and incubated for 24 hr. Neurons were processed for ICC and stained with PHP1 antibody. Part C is similar fractions as in A from different cohorts of mice treated with proteinase K (PK) or untreated and processed similar as in part B. Images were captured with a confocal microscope. (D) Quantification of seeded neurons with fragmented nuclei in each condition is shown. Data show mean ± SEM; ***P<0.001.

## 4. Discussion

This study was designed to investigate the seeding properties of extracellular HTTex1 assemblies in cell models. The major findings were preferential neural entry of seeding-competent HTTex1 species in the sonicated fibrils, nuclear localization, amplification, and induction of neurotoxicity exemplified by depletion of HTT, nuclear damage and caspase-3 activation (Fig. 7). Our studies also implicate the N17 and a prominent epitope within the PRD of HTTex1 regulating the neural entry and amplification of HTTex1 seeds. These findings and the notion that similar seeding-competent mutant HTT may exist *in vivo* suggest that neural uptake and amplification of extracellular amyloidogenic HTTex1 assemblies may constitute a pathogenic pathway in HD.

**Figure 7.**
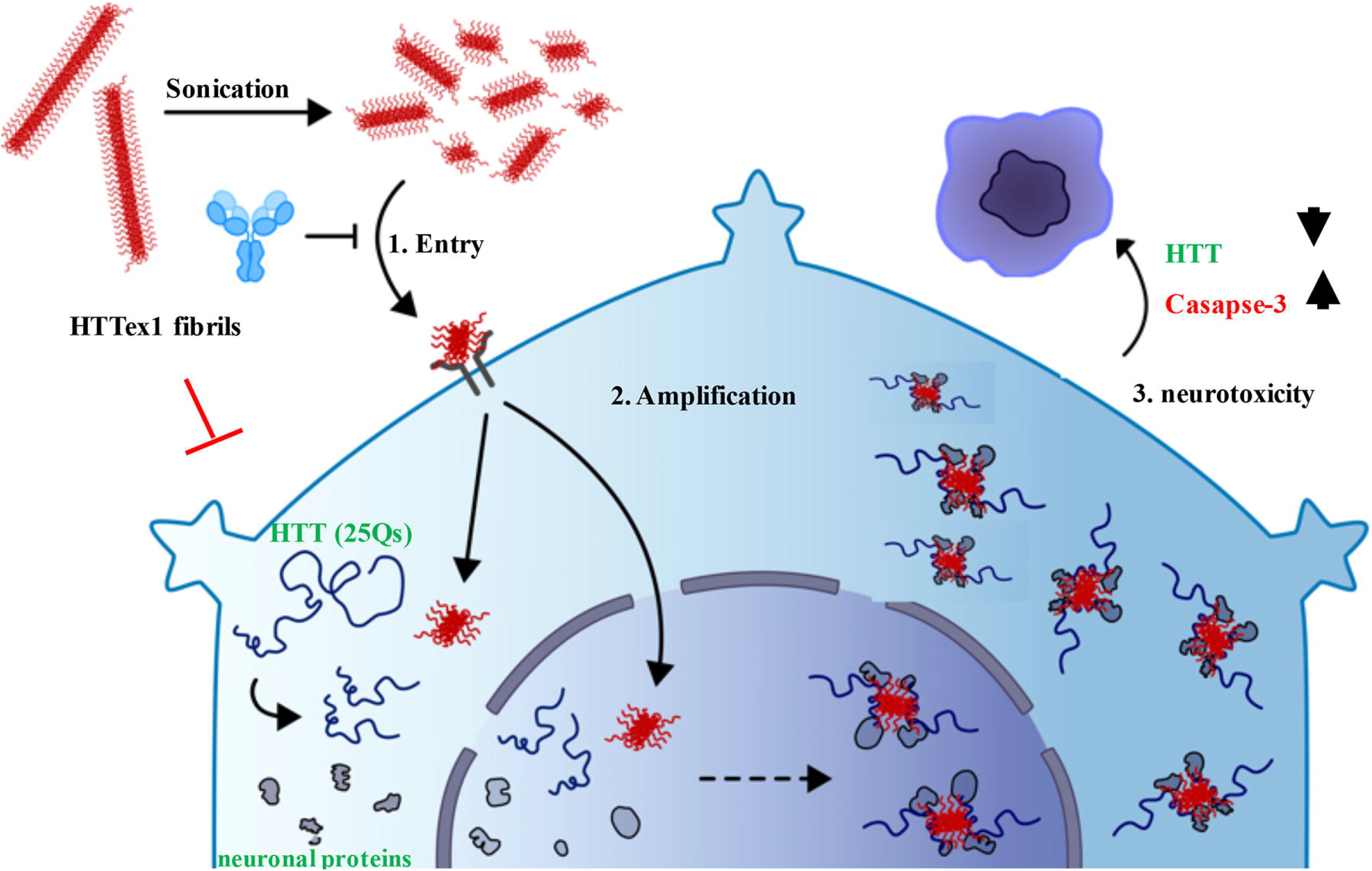
Schematic representation of the extracellular HTTex1 journey in neurons. HTTex1 seeds potentially bind to a surface receptor to enter neurons and ultimately accumulate in the nucleus. Subsequently, the seeds may recruit the endogenous HTT fragments and other neural proteins to amplify. Amplified products induce neurotoxicity potentially by caspase-3 activation and depletion of HTT.

Recent studies suggest that HTTex1 assemblies are taken up by a range of cell lines (Masnata, et al., 2019, Lee, et al. 2020). Our studies indicate that some species in the sonicated HTTex1 fibrils preferentially enter neurons, which is more in line with the neurodegenerative aspect of HD. In our hands, HTTex1 seeds or fibrils did not affect astrocytes or non-neuronal cell lines that were tested. The preference of our HTTex1 seeds for neurons may be due to their structural and biophysical properties imparted by the methods used to prepare the fibrils and/or the evolution of novel conformations induced by sonication. As we are optimizing protocols to assemble and stabilize various oligomeric structures of HTTex1 (Isas, et al., 2019), we may ultimately be able to explore the biological properties of different assemblies of HTTex1 without sonication. This will include structural identity as well as tropism for non-neuronal cells or various neuronal lineages including medium-sized spiny neurons, which appear to degenerate first in HD patients (Bates et al., 2015). The efficient entry and amplification of HTTex1 seeds in our assays is also noteworthy where up to 60% of neurons are affected within 24 hr (Fig. 1A and B). The short incubation period may be due to presence of active receptor(s) on neurons mediating efficient entry and/or the abundance of substrates including the endogenous HTT species for amplification. HTT is abundant in neurons and reduction in the levels of full-length and N-terminal HTT species in the seeded neurons along with their depletion in the lysates seeding assays are consistent with a potential role of endogenous HTT (25Qs) as a substrate for the HTTex1 seeds (Fig. 4). Notably, the HTTex1 seeds initially accumulate in the nuclei of neurons (Fig. 2A), which may be the initial site for amplification. However, since HTT is present in several cellular organelles and due to nuclear-cytoplasmic shuttling properties of HTTex1 (Saudou and Humbert, 2016), multi-focal seeding/amplification may occur. The findings that HTT levels are reduced in the seeded neurons and the amplified products are reactive to several anti-HTT antibodies support a homotypic seeding mechanism (Isas et al., 2017, Ast, et al., 2018, Pandey et al., 2018). Whether heterotypic seeding as demonstrated with α-synuclein and Tau (Jucker and Walker, 2018), also occurs with the HTTex1 seeds in neurons remains to be investigated. However, amyloidogenic HTTex1 species are known to interact with hundreds of cellular proteins, which may also be incorporated into assemblies detected in our assays (Kim et al., 2016, Wanker et al., 2019). The promiscuous interaction of HTTex1 seeds with other neuronal proteins may also contribute to the short incubation period required for assembly formation.

Unraveling the robust neurotoxic aspect of extracellular HTTex1 is another novelty of our work, which might have been difficult to observe previously. Our success might have been a combination of sonicated unbundled HTTex1 fibrils and human neurons with abundant endogenous HTT as several other cell types were resistant to seeds uptake. The pathogenic manifestations of HTTex1 seeds in neurons included nuclear damage, caspase-3 activation and depletion of HTT, which is vital to neuronal survival. Nuclear/DNA damage is detected in the immune cells of prodromal HD patients and is recognized as a pathogenic modifier of HD (Askeland et al., 2018, Castaldo et al., 2019). The mechanism of nuclear damage in our assays remains to be investigated. However, HTTex1 seeds may sequester critical proteins including the endogenous HTT essential for maintaining the nuclear integrity (Kim et al., 2016, Gao et al., 2019, Wanker et al., 2019). HTT is a component of the ATM (ataxia telangiectasia mutated) enzymatic machinery responsible for sensing and repairing DNA damage induced by oxidative stress (Maiuri, et al., 2017). HTT also binds to a complex consisting of RNA polymerase II, CREB binding protein (CBP), ataxin 3, and the DNA repair enzyme polynucleotide-kinase -3- phosphatase (PNKP), which detects and repairs DNA damage generated during transcription elongation (Gao et al., 2019). Thus, depletion of HTT by the HTTex1 seeds may inactivate two prominent sensors of DNA damage and make the nucleus susceptible to insults. The activation of caspase-3 may further contribute to HTTex1-induced nuclear damage and apoptosis. Activated caspase-3 is detected in the brains of HD patients and cleaves HTT in human neurons (Kim et al., 2001, Khoshnan et al., 2009). Moreover, intact HTT is an inhibitor of caspase-3 and reducing its levels promotes cell death (Zhang et al., 2006). While dissecting the precise mechanism of how HTTex1 seeds induce neurotoxicity remain to be characterized, we predict that depletion of endogenous of HTT may impair various physiological pathways in the target neurons, which is manifested as nuclear crumbling and death.

The preference of the HTTex1 seeds for neurons highlights the potential involvement of a cell surface receptor mediating the entry, which is the subject of future studies. On the HTTex1 seeds however, we have found critical epitopes responsible for binding to neuronal surfaces. The antibody inhibition assays and mutagenesis studies indicate that conformations reactive to PHP1/PHP2 antibodies within the PRD (Ko et al., 2018) are important for the neural entry and amplification of HTTex1 seeds. The inability of other PRD-reactive monoclonal antibodies (PHP7-PHP9) to block neural seeding (Fig. 5, Suppl. Fig. 3) reinforces the specificity of novel conformations. Although antibodies to N17 domain also prevented the entry of HTTex1, the seed-antibody complexes remained avidly attached to neurons (Fig. 5, Supplementary fig. 3). We predict that N17 domain is not essential for neuronal binding but the seed-antibody complexes are too large to enter. Alternatively, the N17 domain may be required for events downstream of binding such as transport across neuronal membranes given its high affinity for binding to lipid bilayers (Tao et al., 2019). Recent studies indicate that phosphorylation of residues in the N17 or removal of N17 enhances the neural uptake and nuclear accumulation of HTTex1 fibrils, which also induce neuronal death (Vieweg, et al., 2021). Thus, N17 may regulate some aspect of HTTex1 neuronal entry; albeit serving an inhibitory function. On the other hand, a recent study also found that the anti-N17 antibodies did not prevent the mutant HTTex1 seeding of engineered HEK-293 cells expressing HTTex-1-EGFP cells (Lee et al., 2020). One reason for the discrepancy with our findings may be the variations in the conformations in the N17 to which different antibodies bind. Alternately, the role of N17 in the cell entry of HTTex1 seeds may vary in different paradigms, i.e. neurons versus epithelial cells. Thus, further studies are needed to verify the role of N17 in the neural entry of HTTex1 seeds. Our data however, are consistent with the role of PRD in the HTTex1 seeding HEK-293 cells expressing HTTex1-EGFP (Lee, et al., 2020). Thus, conformations binding the PHP1 and PHP2 antibodies are candidate pathogenic epitopes and may be useful targets to develop reagents for preventing neural seeding *in vivo*. It is encouraging that engineered neurons secreting recombinant PHP2 antibodies are immune to HTTex1 seeds. This may serve as a platform to test other recombinant antibodies for seeding inhibition and ultimately deliver full-length recombinant antibodies to HD models for therapeutic evaluation.

In conclusion, our studies reveal that extracellular seeding-competent HTTex1 may preferentially seed neurons and amplify. To our knowledge this is the first robust neuronal seeding assay where neurotoxic assemblies form by depletion of the normal endogenous HTT. We have further identified the structural determinants of HTTex1 essential for neural seeding. The neuronal assays and reagents including the inhibitory antibodies provide a useful platform to further characterize the biological properties of different HTTex1 seeds, monitor disease progression in HD using fluids such as plasma or CSF, and to further develop compounds, which may prevent the spread and amplification of neurotoxic species in HD. The potential existence of novel receptors for the neural entry of HTTex1 species, which is being investigated by our labs, may further introduce novel pathogenic pathways and therapeutic targets for HD.

## Abbreviations

HD: Huntington’s disease
HTT: huntingtin
HTTex1: huntingtin exon1
polyQ: polyglutamine
PRD: proline-rich domain
CSF: cerebrospinal fluids
IPSC: induced pluripotent stem cells
NPCs: neuronal progenitor cells
WB: Western blots
SDD-AGE: semi-denaturing detergent agarose gel electrophoresis

## Ethics approval

All animal experiments and care complied with federal regulations and were reviewed and approved by the California Institute of Technology Institutional Animal Care and Use Committee (IACUC), protocol#1776-19.

## Funding

Funding was provided by a CHDI award to AK (A-12890), CHDI award (A-12640) to RL and AK, and 1R01NS084345 to RL.

## Authors’ contributions

AK conceived the idea and wrote the manuscript. AC, JMI, NKP, AR, TM performed the experiments. AK, AC, RL and JMI analyzed the data. RL edited the manuscript. All authors have read and approved the paper.

## Declaration of Competing Interest

Authors declare no competing interest.

## Acknowledgements

We are grateful to Dr. Dr. Alejandro Balazs at the Ragon Institute for providing the IgG2 cDNA backbone and Dr. Alysson Muotri at USCD for the IPSC neuronal progenitors. We are also thankful to Dr. Jenny Morton at the University of Cambridge, Dr. Ignacio Munoz-Sanjuan and Dr. Jonathan Bard at CHDI, and Dr. Jeannie Chen at USC for critical suggestions.

## Notes

### Competing Interest Statement

The authors have declared no competing interest.

