## Supplementary figures and images for "Sonicated fibrils of huntingtin exon-1 preferentially seed neurons and produce toxic assemblies"

### Supplemental fig 3

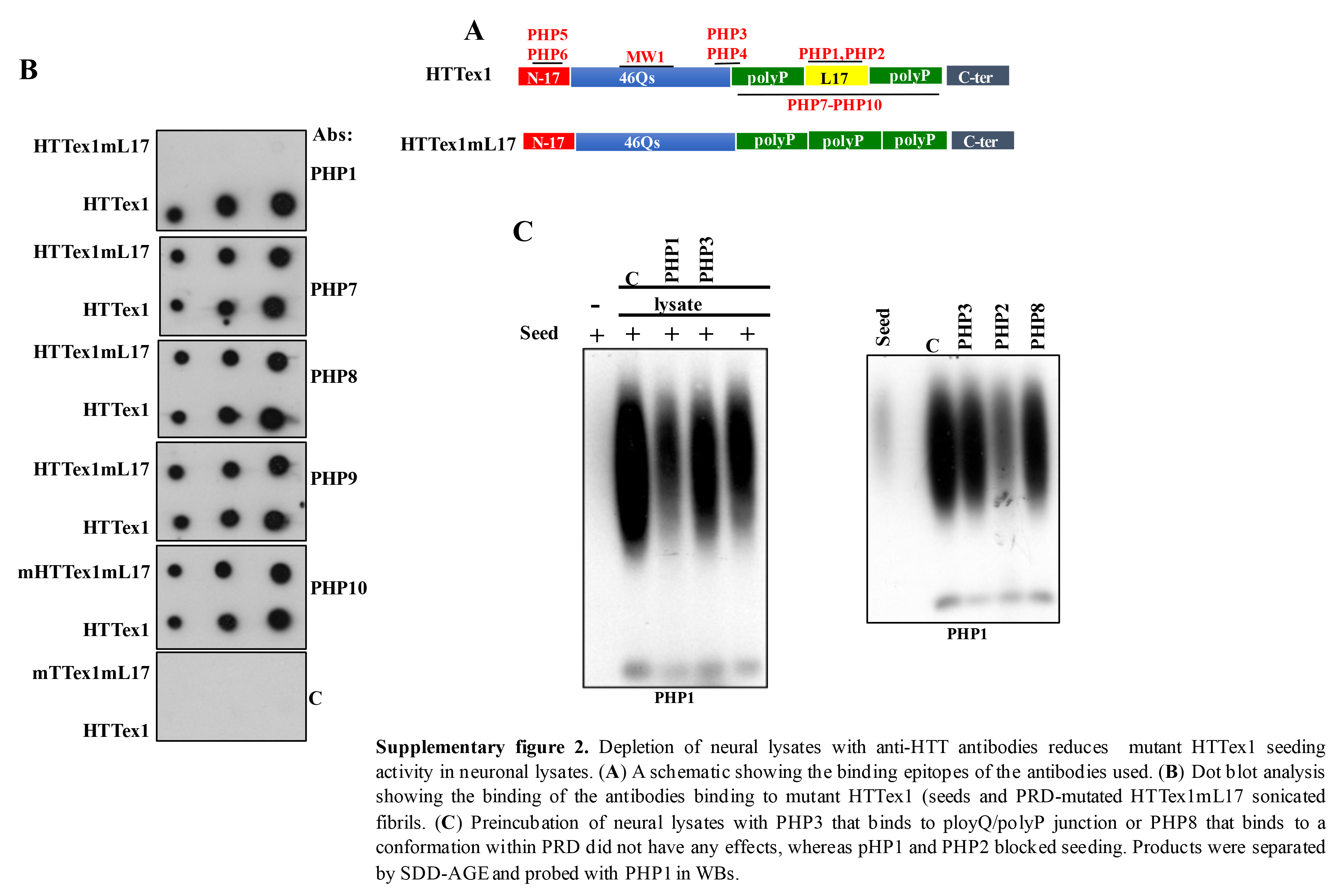

### Supplemental fig. 1

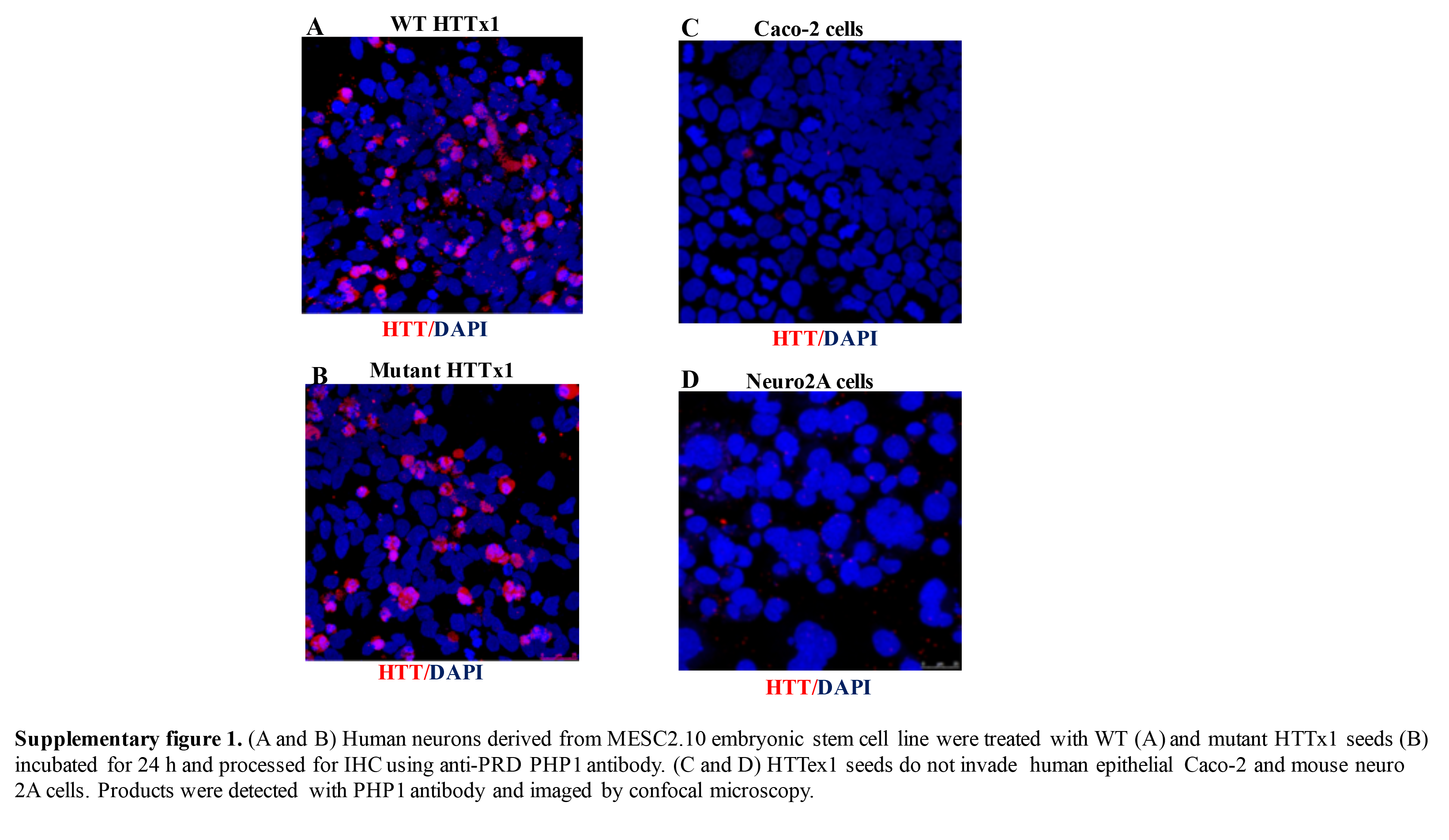

### Supplemental fig. 2

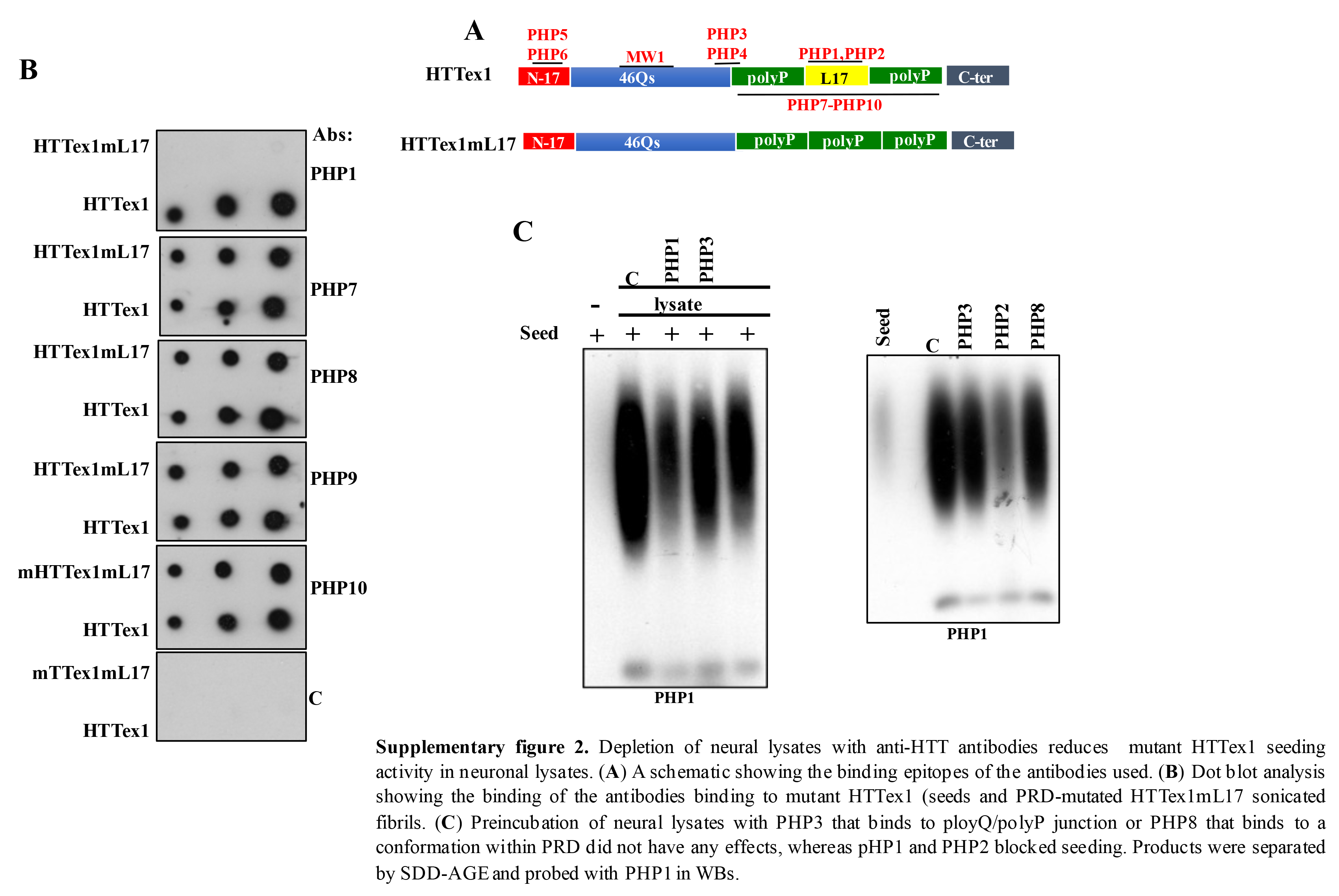

### Supplemental fig. 4

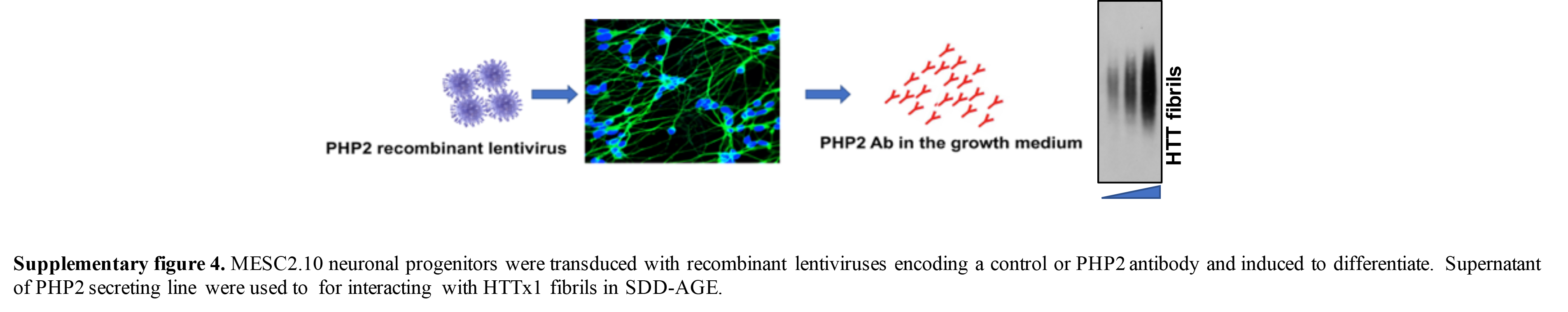
